# Abundant Glomerular Neutrophil Extracellular Traps in C3 Glomerulopathy

**DOI:** 10.64898/2026.08.27.747386

**Authors:** Kim M. O’Sullivan, Priyanka Khandelwal, Patrick D. Walker, Michael J. Hickey, Christoph Licht

**Affiliations:** Centre for Inflammatory Diseases, Monash University Department of Medicine, Monash Medical Centre, Clayton, Victoria, Australia; Division of Nephrology, The Hospital for Sick Children, Toronto, ON, Canada; Department of Paediatrics, University of Toronto, Toronto, ON, Canada; Arkana Laboratories, Little Rock, Arkansas, USA; Cell and Systems Biology Program, Research Institute, The Hospital for Sick Children, Toronto, ON, Canada

**Keywords:** C3 glomerulopathy, proteinuria, neutrophils, neutrophil extracellular traps, complement

## Abstract

**Introduction:** C3 glomerulopathy (C3G) is driven by fluid-phase alternative complement pathway dysregulation, with emerging evidence linking glomerular neutrophil infiltration to disease severity. Neutrophil extracellular traps (NETs) are implicated in other forms of glomerulonephritis. However, their participation in the pathogenesis of C3G remains undefined.

**Methods:** Kidney biopsies from 33 patients with C3G (15 with dense deposit disease [DDD] and 18 with C3 glomerulonephritis [C3GN]) were compared with 15 anti-neutrophil cytoplasmic antibody associated vasculitis (AAV) biopsies as a neutrophil-rich disease control in this retrospective cross-sectional study. Glomerular neutrophils and NETs were identified using immunofluorescence, staining for myeloperoxidase, citrullinated histone H3, peptidyl arginine deiminase-4, and DNA. Supervised machine learning was used to quantify glomerular NET formation, and the data were correlated with kidney function at time of biopsy using linear regression.

**Results:** Intraglomerular NETs were abundant and detected in the majority of glomeruli in C3G biopsies. Compared with AAV, C3G showed a significantly higher fraction of neutrophils forming NETs, despite similar neutrophil counts per glomerulus. NET abundance was similar in DDD and C3GN. In exploratory analyses, a greater proportion of glomeruli containing NETs was associated with lower kidney function (estimated glomerular filtration rate) at biopsy, and this association remained significant after adjustment for age, C3G subtype, and interstitial fibrosis.

**Conclusions:** These observations demonstrate that intraglomerular NETs are a common and prominent observation in C3G and are associated with reduced kidney function at biopsy. These findings raise the possibility that NET deposition in glomeruli is a previously unrecognized driver of glomerular injury in C3G.

**TRANSLATIONAL STATEMENT:** Targeted complement inhibitors have expanded treatment options for C3 glomerulopathy, but treatment responses remain incomplete and the cellular processes accompanying active intrarenal injury remain incompletely defined. We examined renal biopsies of C3G patients for neutrophil infiltration / recruitment and release of neutrophil extracellular traps (NETs) and found that these structures were highly abundant in glomeruli. Greater glomerular NET burden was associated with lower estimated glomerular filtration rate at biopsy. Our observations identify NETs as a candidate component of the pathogenesis of glomerular pathology in C3G and support the investigation of NET-targeting interventions in addition to complement inhibition in C3G patients.

## INTRODUCTION

C3 glomerulopathy (C3G) is a rare, progressive kidney disease driven by dysregulation of the alternative complement pathway.^1, 2^ C3G encompasses C3 glomerulonephritis (C3GN) and dense deposit disease (DDD) and is defined by dominant glomerular C3 deposition.^1, 2^ C3G affects patients across the age spectrum, particularly adolescents and young adults, and presents with proteinuria, hematuria, hypertension, progressive decline in kidney function, and frequent post-transplant recurrence.^3–6^ Prognosis has historically been poor, with up to 50% of adults progressing to kidney failure within 10 years.^7, 8^

Pivotal randomized trials of the C3/C3b inhibitor pegcetacoplan and factor B inhibitor iptacopan demonstrated reductions in proteinuria and stabilization of kidney function, supporting their regulatory approval.^9, 10^ However, despite significant treatment benefits, approximately 30% of patients have persistent proteinuria suggesting previously unrecognized mechanisms not fully addressed by complement inhibition.^9–11^

A growing body of evidence, including abundant neutrophils within C3G glomeruli, the association of a neutrophil-rich glomerular phenotype with complement activation, and enrichment of transcriptional pathways associated with neutrophil recruitment and activation in kidney biopsies, implicates the complement–neutrophil axis in glomerular injury.^12,13^ This concept aligns with observations in other neutrophil-rich glomerular diseases, including systemic lupus erythematosus and ANCA-associated vasculitis (AAV), where activated neutrophils generate NETs, leading to release and deposition of proinflammatory enzymes, proteins, and peptides in the glomerulus.^14–16^ However, the role of intraglomerular neutrophil infiltration and NETs in C3G remains undefined. Therefore, in this study we examined the presence and relevance of neutrophils and NETs in kidney biopsies from patients with C3G.

## SHORT METHODS

Glomerular neutrophil infiltration was quantified via confocal microscopy as number of neutrophils positive for myeloperoxidase (MPO) per glomerular cross section. NETs were assessed by staining for MPO, citrullinated histone H3 (CitH3), peptidyl arginine deiminase-4 (PAD4) and DNA/DAPI. In all biopsies, globally sclerotic glomeruli were excluded from analysis. NET deposition was quantitated using an unbiased automated ImageJ-based algorithm in at least 8 glomeruli per biopsy and expressed as arbitrary units (AU)/glomerular cross section as per previous publications.^16–18^ Between-group comparisons and complete-case linear regression analyses of estimated glomerular filtration rate were performed as described in the **Supplementary Methods**. Baseline patient characteristics were examined using univariate linear regression; variables associated with eGFR at *P* <0.05 were examined in multivariable models.

## RESULTS

### Cohort characteristics

The patient cohort comprised of biopsies examined by Arkana Laboratories from 2016-2022 (with institutional ethics approval IRB 0047). The cohort included 33 patients with C3G (15 DDD and 18 C3GN), 15 patients with AAV, representing a known NET-driven disease, and 6 patients with minimal change disease (MCD) serving as controls. In the C3G cohort, 26 of 33 biopsies were obtained pre-treatment. Baseline clinical and histopathologic features are shown in **Table 1**.

**Table 1.** Baseline patient characteristics at biopsy.

| Characteristic | DDD<sup>a</sup><br>n = 15 | C3GN<sup>b</sup><br>n = 18 | C3G<sup>c</sup><br>n = 33 | AAV<sup>d</sup><br>n = 15 | P<sup>a,b</sup> | P<sup>c,d</sup> |
| --- | --- | --- | --- | --- | --- | --- |
| Age, years | 15.0 (7.0 - 20.0) | 24.0 (16.0 - 33.0) | 18.0 (13.0 - 29.0) | 64.0 (42.0 - 71.0) | **0.03** | **0.001** |
| Female | 12 (80.0) | 11 (61.1) | 23 (69.7) | 9 (60.0) | 0.3 | 0.5 |
| Proteinuria, g/day or g/g | 3.5 (2.5 - 15.0) | 5.6 (1.0 - 6.9) | 4.6 (2.5 - 8.0) | 2.0 (1.9 - 3.0) | 0.6 | 0.2 |
| Missing | 8 | 11 | 19 | 10 |  |  |
| Proteinuria |  |  |  |  | 0.3 | 0.2 |
| Nephrotic | 11 (91.7) | 9 (69.2) | 20 (80.0) | 5 (55.6) |  |  |
| Sub-nephrotic | 1 (8.3) | 4 (30.8) | 5 (20.0) | 4 (44.4) |  |  |
| Missing | 3 | 5 | 8 | 6 |  |  |
| Hematuria, gross/microscopic | 8 (88.9) | 6 (100.0) | 14 (93.3) | 11 (100.0) | 1.0 | 1.0 |
| Missing | 6 | 12 | 18 | 4 |  |  |
| eGFR, mL/min/1.73 m<sup>2</sup> | 90.0 (77.6 - 127.8) | 82.7 (9.3 - 100.1) | 83.2 (37.0 - 104.4) | 20.5 (8.3 - 23.4) | 0.1 | **0.003** |
| Missing | 6 | 3 | 9 | 4 |  |  |
| eGFR < 90 mL/min/1.73 m<sup>2</sup> | 4 (44.4) | 7 (46.7) | 11 (45.8) | 11 (100.0) | 1.0 | **0.002** |
| Missing | 6 | 3 | 9 | 4 |  |  |
| Low C3 |  |  |  |  | **0.03** | 0.1 |
| Normal | 0 (0.0) | 4 (44.4) | 4 (21.1) | 3 (60.0) |  |  |
| Low | 10 (100.0) | 5 (55.6) | 15 (78.9) | 2 (40.0) |  |  |
| Missing | 5 | 9 | 14 | 10 |  |  |
| Low C4 |  |  |  |  | 0.6 | 1.0 |
| Normal | 6 (66.7) | 5 (83.3) | 11 (73.3) | 4 (80.0) |  |  |
| Low | 3 (33.3) | 1 (16.7) | 4 (26.7) | 1 (20.0) |  |  |
| Missing | 6 | 12 | 18 | 10 |  |  |
| Treatment naive | 14 (93.3) | 12 (66.7) | 26 (78.8) | N/A |  |  |
| Number of glomeruli/patient | 18.0 (13.0 - 36.0) | 23.5 (18.0 - 45.0) | 20.0 (16.0 - 36.0) | 17.0 (12.0 - 26.0) | 0.4 | 0.2 |
| C3HI Activity Index (out of 21) | 7.0 (3.0 - 8.0) | 4.0 (3.0 - 9.0) | 5.0 (3.0 - 8.0) | n/a | 0.7 |  |
| C3HI Chronicity Index (out of 10) | 0.0 (0.0 - 2.0) | 5.0 (2.0 - 7.0) | 2.0 (0.0 - 6.0) | n/a | **0.002** |  |
| Number of neutrophils per glomerular cross-section | 10.8 (5.0 - 18.8) | 11.0 (6.5 - 26.0) | 10.8 (5.5 - 20.5) | 14.8 (11.0 - 22.0) | 0.4 | 0.2 |
| NETs per glomerular cross-section | 5.5 (2.9 - 10.8) | 7.5 (3.3 - 12.1) | 6.0 (3.3 - 10.8) | 7.5 (3.0 - 10.3) | 0.4 | 0.9 |
| Glomerular area with NETs | 4.3 (0.5 - 6.9) | 5.4 (3.5 - 9.5) | 5.0 (3.0 - 8.0) | 3.2 (1.8 - 7.1) | 0.1 | 0.6 |
| Percent glomeruli with NETs | 87.0 (33.0 - 100.0) | 100.0 (100.0 - 100.0) | 100.0 (83.0 - 100.0) | 62.5 (50.0 - 100.0) | 0.1 | 0.05 |
| Percent neutrophils with NETs | 53.5 (32.0 - 68.0) | 60.2 (49.5 - 71.0) | 54.0 (45.4 - 69.1) | 33.9 (19.6 - 50.0) | 0.3 | **0.005** |

Values are presented as median (interquartile range) or number (percentage).
DDD, dense deposit disease; C3GN, C3 glomerulonephritis; C3G, C3 glomerulopathy; AAV, antineutrophil cytoplasmic antibody associated vasculitis; eGFR, estimated glomerular filtration rate; C3HI, C3 glomerulopathy histologic index; NETs, neutrophil extracellular traps. P<sup>a,b</sup> denotes comparison between DDD and C3GN; P<sup>c,d</sup> denotes comparison between C3G and AAV

### Glomerular NETs are abundant in C3G

Glomerular NETs, defined by colocalization of extracellular DNA with MPO, CitH3, PAD4, and DNA/DAPI, were detected across the C3G cohort, with a median of 100% of glomeruli containing NETs (IQR, 83-100; **Figure 1 & Table 1**). In patients with DDD **(Figure 1A)** and C3GN **(Figure 1B)**, glomerular neutrophils, as highlighted by MPO staining, were abundant and located throughout glomeruli, while NETs were particularly enriched in areas of hypercellularity with neutrophilic accumulation. In contrast, NET formation was rarely observed in the tubulointerstitium of the C3G cohort highlighting glomerulus-specific retention of neutrophils and NET formation. To gauge the relative extent of NET deposition in the C3G cohort, neutrophil and NET frequency were compared with that in the established NET-driven disease of AAV **(Figure 1C)**. Relative to AAV, C3G showed a significantly higher proportion of glomerular neutrophils undergoing NET formation (54.0% [IQR, 45.4–69.1] versus 33.9% [IQR, 19.6–50.0], *P* = 0.005). The percentage of glomeruli containing NETs was also higher in C3G than in AAV (100% [IQR, 83–100] versus 62.5% [IQR, 50–100], *P* = 0.05). In contrast, the number of neutrophils per glomerulus was similar between C3G and AAV. Together, these data suggest that neutrophils recruited to glomeruli of C3G patients have a higher propensity to NET than those in AAV patients.

**Figure 1.**
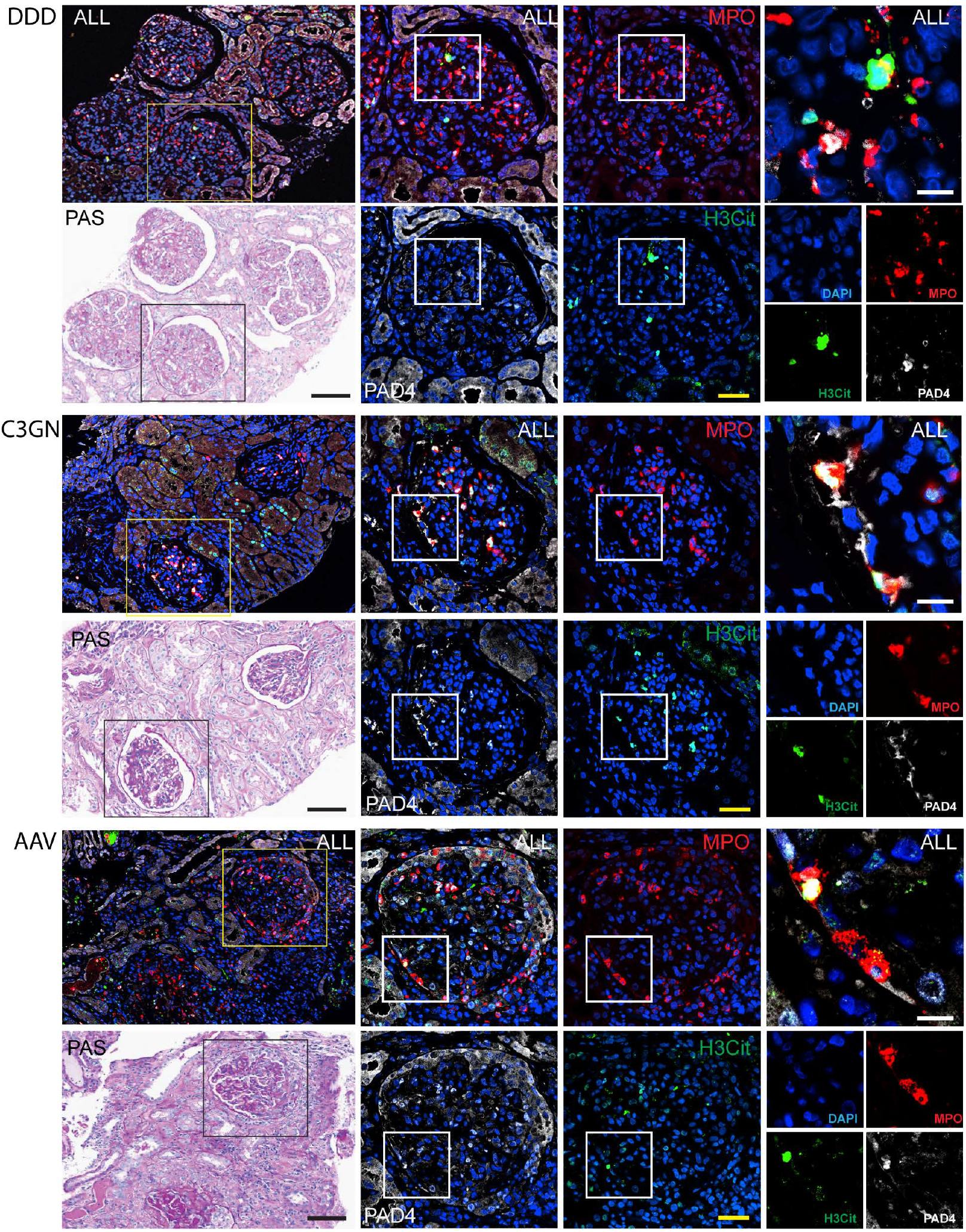
NET deposition and renal pathology in DDD, C3GN and AAV kidney biopsies. (a) DDD, NET deposition, identified as MPO (red), PAD4 (grey), H3Cit (green) and DNA (blue) and PAS staining showing segmental necrosis and mesangial thickening. (b) C3GN NET deposition, identified as MPO (red), PAD4 (grey), H3Cit (green) and DNA (blue) and PAS staining showing segmental necrosis and mesangial thickening. (c) AAV, NET deposition, identified as MPO (red), PAD4 (grey), H3Cit (green) and DNA (blue) and PAS staining showing segmental necrosis and crescent formation. Original magnification 200x (low power) and 600x (high power). Scale bars denote 100 µm (black), 50 µm (yellow) and 10 µm (white).

### Greater NET burden is associated with impaired kidney function

To examine the clinicopathologic relevance of NETs in C3G, we evaluated the associations of kidney function (eGFR) at biopsy with both the frequency and extent of glomerular NET deposition. A greater proportion of glomeruli containing NETs was associated with lower eGFR. A 10% increase in glomeruli containing NETs was associated with a decrease in eGFR of 9.5 mL/min/1.73 m^2^ (95% CI, −15.8 to −3.2; *P* = 0.005; **Table 2**). The association remained significant in a multivariable model after adjustment for age, C3G subtype, and interstitial fibrosis (−8.9 mL/min/1.73 m^2^; 95% CI, −14.0 to −3.7; *P* < 0.01; **Table 2**). The proportion of neutrophils forming NETs was similarly associated with lower eGFR in unadjusted analysis (−11.1; 95% CI, −20.1 to −2.2; *P* <0.05; **Table 2**), but the association was attenuated after adjustment (−7.6; 95% CI, −16.2 to 0.9; *P* = 0.08; **Table 2**).

**Table 2.** Glomerular NET measures associate with eGFR in C3G.

| Parameter | Univariable model, $\beta$<br>(95% CI) | *P*<br>value | Multivariable:<br>Glomeruli with NETs<br>(N=24), $\beta$ (95% CI) | *P*<br>value | Multivariable: Neutrophils<br>with NETs,<br>$\beta$ (95% CI) | *P*<br>value |
| --- | --- | --- | --- | --- | --- | --- |
| Age, per 1-year increase | -1.7 (-2.6 to -0.7) | **0.001** | -1.4 (-2.3 to -0.5) | **0.006** | -1.3 (-2.4 to -0.1) | **0.03** |
| Male versus female | -27.4 (-76.1 to 21.2) | 0.25 | — | — | — | — |
| C3GN versus DDD | -49.2 (-92.9 to -5.5) | **0.03** | 6.9 (-32.8 to 46.7) | 0.72 | -5.8 (-51.9 to 40.3) | 0.79 |
| Low versus normal serum C3 | 54.9 (-8.7 to 118.4) | 0.09 | — | — | — | — |
| Low versus normal serum C4 | -35.8 (-183.7 to 112.0) | 0.60 | — | — | — | — |
| Crescents, per 10-percentage-point increase | -2.4 (-10.3 to 5.5) | 0.54 | — | — | — | — |
| Mesangial hypercellularity, per one-grade increase | 16.8 (-4.3 to 37.8) | 0.11 | — | — | — | — |
| Endocapillary hypercellularity, per one-grade increase | 13.2 (-6.2 to 32.7) | 0.17 | — | — | — | — |
| Interstitial inflammation, per one-grade increase | -13.8 (-42.2 to 14.7) | 0.33 | — | — | — | — |
| Interstitial fibrosis, per one-grade increase | -28.3 (-46.9 to -9.7) | **0.005** | -9.2 (-28.7 to 10.4) | 0.34 | -6.4 (-30.3 to 17.6) | 0.59 |
| Glomeruli with NETs, per 10-percentage-point increase | -9.5 (-15.8 to -3.2) | **0.005** | -8.9 (-14.0 to -3.7) | **0.002** | — | — |
| Neutrophils with NETs, per 10-percentage-point increase | -11.1 (-20.1 to -2.2) | **0.02** | — | — | -7.6 (-16.2 to 0.9) | 0.08 |

CI, confidence interval; C3G, C3 glomerulopathy; C3GN, C3 glomerulonephritis; DDD, dense deposit disease; eGFR, estimated glomerular filtration rate; NETs, neutrophil extracellular traps. Female sex, DDD, and normal serum C3 and C4 were the reference categories for categorical variables. $\beta$ coefficients for categorical variables represent the estimated difference in eGFR relative to the reference category. $\beta$ coefficients for NET measures represent the estimated change in eGFR, in mL/min/1.73 m<sup>2</sup>, per 10-percentage-point increase. Age coefficients represent the estimated change in eGFR per 1-year increase. Interstitial fibrosis was coded as 0, none; 1, 1%-25%; 2, 26%-50%; and 3, >50% and modeled as linear score from 0-3. The interstitial fibrosis coefficient represents the estimated difference in eGFR per one-category increase. Both multivariable models included 24 patients with complete data for eGFR, age, C3G subtype, interstitial fibrosis, and the respective NET measure.

Neither the proportion of glomeruli containing NETs nor the proportion of neutrophils forming NETs was associated with the C3G histologic activity index or chronicity index (data not shown).

## DISCUSSION

This study identifies glomerular neutrophils and NET formation as an important component in the pathology of C3G. C3GN and DDD showed similar neutrophil and NET deposition, indicating that NETs are a common feature across the C3G spectrum. Despite comparable glomerular neutrophil counts, C3G biopsies contained a greater proportion of NET-forming neutrophils and NET-containing glomeruli than AAV biopsies. The comparison with AAV is informative, since in AAV, NET deposition is an established mechanism of glomerular injury.^16, 18–20^ The current findings therefore place C3G within a broader group of inflammatory glomerular diseases, in which NETs are associated with glomerular injury.

Previous biopsy-based studies in C3G have shown that glomerular C5b-9 deposition is associated with greater histological chronicity and adverse kidney outcomes.^13^ The biopsy cluster characterised by high glomerular C5b-9 and worse kidney survival showed a distinct immune cell- and fibroblast-associated signature, with enrichment of transcriptional pathways involving leukocyte adhesion, activation, and survival.^13^ A subsequent spatial transcriptomics analysis of C3G biopsies demonstrated that neutrophil- and macrophage-rich glomerular inflammation was associated with intrarenal complement activation and proteinuria.^12^ Our findings extend this framework by showing that glomerular neutrophils in C3G are activated toward NET formation.

Histopathologically, NETs were frequently enriched in areas of glomerular damage. Moreover, a greater proportion of glomeruli containing NETs was associated with impaired kidney function, and this effect was independent of age, C3G subtype and interstitial fibrosis. The proportion of NET-forming neutrophils showed a similar inverse association with kidney function in the univariate analysis. Together, these findings are complementary evidence of a functional role of glomerular NET formation in C3G pathology and, as kidney function at biopsy is known to predict progression to kidney failure, may serve as novel prognostic marker.^21, 22^ Given the small sample size, these findings need validation in larger longitudinal cohorts.

The literature holds evidence for neutrophils, NETs and complement forming a vicious cycle in which complement can prime neutrophils (via C3b and C5a) and promote NETosis, and NETs can act as scaffold for complement deposition and further amplification.^23, 24^ Currently available complement inhibitors may reduce neutrophil priming, activation and recruitment. However, we speculate that in patients less responsive to these treatments, NET formation may be ongoing and potentially be amplified by alternative NET triggers such as cytokines, infectious agents and immune complexes.^25^ Such patients might benefit from adjunctive NET-targeted therapies alongside new generation complement inhibitors.

### Limitations

The present study should be viewed as hypothesis-generating: the sample size is modest (with incomplete availability of some clinical and complement assessments), analysis is cross-sectional, and causality cannot be inferred with certainty. Prospective studies integrating evaluation of kidney tissue with longitudinal clinical outcomes, will be important to determine whether glomerular NET formation is a marker of active injury, a mediator of progression, or both.

## Conclusions

These findings identify glomerular neutrophil infiltration and NET formation as so far unrecognized prominent features of C3G that are associated with lower kidney function at biopsy and raise the possibility that they represent a novel driver of glomerular injury in C3G. This complement-neutrophil-NETs axis provides a rationale for future mechanistic and therapeutic studies evaluating the potential of NET targeting treatments to improve patient outcomes.

## ACKNOWLEDGEMENTS FUNDING STATEMENT

This work was supported by CIHR Project Grant #542895 (NPA Christoph Licht). The authors acknowledge Monash Micro Imaging at the Monash Health Translation Precinct (MMI-MHTP) for the provision of microscopy equipment and the Monash Histology Platform for microtomy of biopsies. The authors acknowledge Arkana Laboratories for curation of patient biopsies.

## DISCLOSURES / CONFLICTS OF INTEREST

KMOS reports research funding via consultancy and grants from Patrys Ltd. PW & MJH declare no conflicts of interest. CL is an advisory board member and invited speaker for Alexion, AstraZeneca, Novartis, and Sobi.

## DATA SHARING STATEMENT

Data is available from the Corresponding Authors on reasonable request.

## AUTHOR CONTRIBUTIONS

KMOS, MJH and CL conceptualized the study. KMOS performed all experimental work, image acquisition and analysis. PDW assessed pathology of biopsies, PK performed clinical correlation analysis. KMOS, PK, PDW, MJH and CL wrote the original manuscript.

