## Supplemental File for "Abundant Glomerular Neutrophil Extracellular Traps in C3 Glomerulopathy"

### Supplemental Methods

#### Study design, patient selection and histological assessment

This retrospective cross-sectional study included archived native-kidney biopsies. The study included patients with C3G, ANCA-associated vasculitis (positive control), and minimal change disease [MCD] (negative control). De-identified slides and reports were examined. Clinical data was gathered from the material provided in the biopsy report. Renal biopsies were processed for light, immunofluorescence and electron microscopy using standard techniques.[1] Renal biopsy criteria for C3GN and DDD required C3 dominant immunofluorescent findings ≥ 2+ (on a 0 to 3+ scale) and two orders of magnitude greater than other immunoreactants. All cases underwent pronase digestion of formalin-fixed paraffin-embedded tissue to rule out masked immunoglobulin deposition, followed by staining for IgG, IgA, IgM and kappa and lambda chains.[2] Electron microscopy was used to further distinguish between C3GN and DDD (dense deposits). EM samples were taken from formalin fixed tissue and processed into plastic blocks, cut with a diamond knife and examined with a JEOL electron microscope.

In all cases biopsy findings were correlated with clinically documented abnormalities of the alternative pathway of complement for final diagnosis. Renal biopsy criteria for AAV included evidence of pauci-immune necrotizing and crescentic glomerulonephritis along with positive ANCA sera with either negative or minimal (pauci) staining for immunoglobulins and complement. Biopsies were eligible to be included when the diagnosis of the above had been established and sufficient tissue was available for quantitative immunofluorescence analysis. Biopsies were excluded if there was inadequate tissue for further studies. For patients with more than one biopsy only the first diagnostic biopsy was used. The study was approved by the Solutions IRB under the Protocol 0047: A Novel Role for Neutrophils and NETs in the

Pathogenesis of C3 Glomerulopathy. Informed consent was waived. C3G histologic activity and chronicity indices were determined as previously described.[3, 4]

**Clinical data and definitions**

Clinical and laboratory data were obtained from records accompanying the diagnostic biopsy. Proteinuria was expressed as 24-hour urine protein excretion in g/day or as a spot urine protein-to-creatinine ratio in g/g. Nephrotic-range proteinuria was defined as >2 g/g in children or >3.5 g/day in adults. Serum C3 and C4 were classified as low when below the lower limit of the reporting laboratory's reference range. eGFR was estimated using the serum creatinine concentration measured closest to kidney biopsy. For patients aged <18 years, eGFR was calculated using the height-independent, creatinine-based Full Age Spectrum (FAS) equation;[5] for patients aged ≥18 years, eGFR was calculated using the race-free 2021 CKD-EPI creatinine equation.[6] Treatment-naïve status was defined as the absence of prior treatment with corticosteroids, other immunosuppression, or complement inhibitors before the index biopsy.

**NET identification and quantification**

Formalin-fixed paraffin-embedded 3-micron thick kidney biopsy sections were adhered to Superfrost Plus slides (Menzel), heated in a 60-degree oven to allow sections to adhere firmly, then dewaxed using xylene, rehydrated in ethanol and washed in phosphate buffered saline (PBS). Antigen retrieval was performed using Tris-EDTA buffer (10mM Tris, 1mM EDTA, pH 9.0) in a pressure cooker for 10 minutes followed by a subsequent 20-minute cool down (as described previously).[7] Sections were washed twice in PBS and blocked with 10% chicken sera in 10% PBS/Bovine Serum Albumin (BSA) for 30 minutes. NETs were identified using the gold standard technique of staining with MPO, PAD4, H3Cit and DAPI (for DNA) as previously described.[7] Primary antibodies were incubated overnight at 4 degrees in a humidity chamber, at the following concentrations: goat anti-human MPO (AF3677, R&D Systems, 10 µg/ml), mouse anti-human PAD4 (ab128086, 10 µg/ml, Abcam), and rabbit anti-human H3Cit (ab5103, 5 µg/ml, Abcam). Mouse IgG (purified in house), Rabbit IgG (purified from whole serum from Sigma) and Donkey IgG (purified from whole serum from Sigma) used at the same concentrations served as isotype controls. The following day the sections were washed in PBS and the secondary antibodies applied for 40 minutes in 1% BSA/PBS as follows: chicken anti goat Alexa Fluor 594, chicken anti rabbit Alexa Fluor 488 and donkey anti mouse Alex Fluor 647 (Molecular Probes, Thermo Fisher Scientific, A-21468, A-21441, and A-31571; all 1:200 in 1%BSA/PBS). Tissues were subsequently washed twice in PBS then incubated with DAPI for 10 minutes and washed. Kidney tissue autofluorescence was quenched using Sudan black (0.1% in 70% ethanol for 30 minutes) prior to coverslipping with anti-fade mounting media ProLong Gold (Molecular probes, Thermo Fisher) and #1.5 glass confocal coverslips (HD Scientific). Images were captured using a Nikon AxR inverted confocal microscope with resonant scanner, spectral imaging and Nikon's "Denoise.ai" algorithm using laser lines 405nm, 488nm, 561nm and 640nm (Nikon, Japan). Single plane images (1024 x 1024 x 16 bit) were captured in a line sequential manner (4 line averaging) using the 20x objective lens for low powered images and the 60x oil objective lens for high powered images.

**Quantification and analysis**

NET formation was analysed using open-source supervised machine learning tools in image analysis software Image J (NIH, Bethseda). The Image J trainable plugin Weka segmentation was utilized to identify the number of glomerular NETs and number of neutrophils forming NETs as previously described.[8-10] This computational model uses classifiers to train the model to recognise NETs as MPO positive cells (neutrophils) that are colocalized with H3Cit, PAD4 and DNA (detected via DAPI positive nuclei and extracellular DNA). Glomeruli were isolated using the region of interest (ROI) tool in Image J, and numbers of NETs and number of neutrophils forming NETs calculated per glomerular cross section.

**Statistical analysis**

Continuous variables were summarized as median (interquartile range) and categorical variables as number (percentage). Between-group comparisons were performed using the Wilcoxon rank-sum test or Fisher's exact test as appropriate. Within the C3G cohort, associations of the percentage of glomeruli containing NETs and neutrophils forming NETs with C3G histologic activity and chronicity indices were examined using separate univariate linear regression models. Further, associations between eGFR at biopsy (dependent variable) and candidate clinical and histopathologic variables - including age, sex, C3G subtype, serum C3 and C4, percentage glomeruli with crescents, mesangial and endocapillary hypercellularity, interstitial inflammation, interstitial fibrosis, percentage of glomeruli containing NETs, and percentage of neutrophils forming NETs - were first examined using univariate linear regression. Variables associated with eGFR at $P < 0.05$ were examined in multivariable models. Each NET measure was evaluated separately in a multivariable model including age, C3G subtype and interstitial fibrosis.

Multivariable analyses included patients with complete data. Beta coefficients represent the change in eGFR, in mL/min/1.73 $m^2$, per 10-percentage-point increase in NET measures. Variance inflation factors were low in all adjusted models (maximum, 2.07), and normal quantile-quantile plots showed no major departures from residual normality. Because of the small sample size, multivariable analyses were considered exploratory and hypothesis-generating. Two-tailed $P < 0.05$ was considered significant. Analyses were performed using R version 4.5.2. (R Core Team, 2025).
